# Molecular Investigation of TSHR gene in Bangladeshi Congenital Hypothyroid patients

**DOI:** 10.1101/2023.02.20.529210

**Authors:** Mst. Noorjahan Begum, Rumana Mahtarin, Md Tarikul Islam, Sinthyia Ahmed, Tasnia Kawsar Konika, Kaiissar Mannoor, Sharif Akhteruzzaman, Firdausi Qadri

**Affiliations:** Institute for Developing Science and Health Initiatives (ideSHi), 167/24, Blue Moon Gram Tower, Kalshi Road, ECB Chattar, Mirpur, Dhaka-1216, Bangladesh; Department of Genetic Engineering & Biotechnology, University of Dhaka, Dhaka-1000, Bangladesh; Department of Biochemistry and Molecular Biology, Shahjalal University of Science and Technology, Sylhet-3114; Division of Computer Aided Drug Design, The Red-Green Research Centre, BICCB, Tejgaon, Dhaka, Bangladesh; Nuclear Medicine and Allied Sciences, Bangabandhu Sheikh Mujib Medical University (BSMMU), Shahbag, Dhaka-1000, Bangladesh; Virology Laboratory, Infectious Diseases Division, International Centre for Diarrhoeal Disease Research, Bangladesh, Mohakhali, Dhaka, Bangladesh; Mucosal Immunology and Vaccinology, Infectious Diseases Division, International Centre for Diarrhoeal Disease Research, Bangladesh, Mohakhali, Dhaka, Bangladesh

**Keywords:** Congenital Hypothyroidism, Thyroid Dysgenesis, TSHR, Sequencing, Molecular docking, Non-covalent interaction, Molecular dynamics, Principle component analysis

## Abstract

The disorder of thyroid gland development or thyroid dysgenesis (TH) accounts for 80-85% cases of congenital hypothyroidism (CH). Hence, the understanding of molecular etiology of TH is prerequisite. Mutations in TSHR gene is mostly associated with thyroid dysgenesis, prevent or disrupt normal development of the gland. The current study detects two nonsynonymous mutations (p.Ser508Leu, p.Asp727Glu) in transmembrane (TM)-region (Exon 10) of TSHR gene in 21 patients with dysgenesis by sequencing-based analysis. Later, transmembrane (TM)-region of TSHR protein is modelled by homology modeling. Transmembrane (TM)-region of TSHR protein is targeted by small molecules thyrogenic drugs, MS437 and MS438 to perceive the effect of mutations. The damaging effect in drug-protein complexes of mutants were envisaged by molecular docking and interactions. The binding affinity of wild type protein was much higher than the mutant cases for both of the ligands (MS437 and MS438). Molecular dynamics simulates dynamic behavior of wild type and mutant complexes. MS437-TSHR_368-764_MT2 and MS438-TSHR_368-764_MT1 show stable conformations in biological environments. Finally, PCA reveals structural and energy profile discrepancies. TSHR_368-764_MT1 exhibits much variations than TSHR_368-764_WT and TSHR_368-764_MT2, emphasizing more damaging pattern in TSHR_368-764_MT1. The study might be helpful to understand molecular etiology of thyroid dysgenesis (TH) exploring the mutational impact on TSHR protein to the interaction with agonists.

## 1. Introduction

Congenital hypothyroidism can be prompted by variable factors in which only some are coupled with genetics. Genetic causes account for about 15 to 20 percent of cases of congenital hypothyroidism. Although CH is a genetically heterogeneous disorder, the candidate genes divide the disorder into two main groups namely thyroid dysgenesis and thyroid dyshormonogenesis. Different studies including online databases such as Genetics Home Reference and Online Mendelian Inheritance in Men (OMIM) suggested that about 10-20 percent of total cases with CH were associated with thyroid dyshormonogenesis that would result from mutations in one of several genes involved in the biosynthesis of thyroid hormones (1). The above-mentioned databases also described that about 80-85% of CH cases are associated with disorders of thyroid gland development (Dysgenesis) which is categorized by ectopic (located in a distant region, 40%), agenesis (absent of thyroid gland, 40%), and the other cases are accompanying with hypoplasia (small size). Although the actual cause of thyroid dysgenesis is still under investigation, some studies have suggested that 4 major genes which play roles in the proper growth and development of the thyroid gland, such as TSHR (Thyroid 3 stimulating hormone receptor) and three transcription factors-TTF-1, TTF-2, and PAX8 (paired box-8, transcription factor) (2) (3). Mutations in these genes prevent or disrupt normal development of the gland. TSHR gene is predominantly related with thyroid dysgenesis, as most of the mutations occurs in the gene in CH patients (4). TSHR is a G protein coupled transmembrane receptor which is present on the surface of thyroid follicular cells. TSH, secreted by the anterior pituitary, mediates its effect through TSHR which is crucial for thyroid gland development and function. TSHR gene is located on chromosome 14q31 and contains 11 Exons code for a receptor protein of 764 amino acid residues (5, 6). TSHR has high affinity binding sites for TSH. Mutations in TSHR gene results in mutant TSHR protein which lacks its binding affinity to TSH or loses its ability to activate adenylate cyclase. And thus, mutant TSHR protein disrupts thyroid gland development and proper functioning. TSHR mutation may also be present with a normally placed thyroid gland. TSHR gene mutation is reported to be inherited as autosomal recessive manner and Exon 10 is known to carry majority of the mutations (7). Previously two small molecule agonists (MS437 and MS438) with the highest potency bind to the transmembrane domain of the TSHR receptor and initiated signal transduction by activating the transmembrane domain. These small molecules (MS437 and MS438) also showed TSHR gene expression. (8) Hence, in the present study we have explored the effect of our detected mutations in the 3D structure of TSHR transmembrane (TM)-region targeted by thyrogenic drugs, MS437 and MS438.

## 2. Methods and Materials

### 2.1. Study Design, Clinical Settings and Ethical Clearance

The study was designed carried out on 21 confirmed cases of Congenital Hypothyroid children who are kept under treatment of Levothyroxine (LT4) drug.in the Department of Endocrinology and National Institute of Nuclear Medicine and Allied Sciences (NINMAS) of Bangabandhu Shaikh Mujib Medical University (BSMMU). Ethical permission was obtained from the Ethical Review Committee of University of Dhaka (CP-4029) and the study was collaborated with NINMAS and Dept. of Endocrinology, BSMMU for specimen collection. Prior to enrollment of study participants, a written informed consent along with the clinical information was collected from the parent(s) or legal guardian(s) of each patient.

### 2.2. Collection and Processing of Blood Specimens

Blood Specimens were collected from the participants to conduct the molecular, biochemical and metabolic profiling tests. A total of 5ml blood was collected from each participant. Then it was aliquoted in different tubes for different types of analysis, such as (1) 3 mL blood in EDTA tube for DNA isolation, (2) 75 µl blood to prepare Dried Blood Spots (DBS, Whatman® 903 generic multipart filter paper, GE Healthcare, Westborough, USA) for metabolic profiling, (3) 250 µl of blood in tube containing 750 µl Trizol® LS (Ambion, Life Technology, Cat. No. 10296-028) for conducting genetic study using RNA and (4) rest of the blood was used for separation of plasma-serum for biochemical test. All the samples were transported to the laboratory immediately. The DBS cards were dried for at least 4 hours at room temperature and once dried, they were stored at −70°C freezer with desiccant for protection from moisture. All the plasma, serum and Trizol containing specimens 17 were stored at −20°C freezer. After the genomic DNA isolation, EDTA containing blood was stored at −70°C freezer.

### 2.3. Molecular Analysis of TSHR Gene

Now-a-days, gene-based study plays the key role to explore the actual cause of a particular disease. The present study was designed to perform the molecular analysis in various steps.

#### 2.3.1. Genomic DNA Isolation to Perform PCR

Genomic DNA was isolated from the EDTA blood by using Qiagen DNAeasy mini kit according to the manufacturer’s instruction. 500 µl of FG1 buffer was taken in a 1.5 ml microcentrifuge tube. 200 µl of whole blood was added to the FG1 buffer and mixed by inverting the tube 5-10 times. The mixture was then centrifuged at 10,000 x g for 5 minutes in fixed angle rotor. The supernatant was carefully removed so that the pellet remained in the tube. 1µl of QIAGEN protease was added to 100 µl of FG2 buffer and mixed by vortex in a fresh Eppendorf tube. Then 100 µL of FG2/QIAGEN protease was added to the pellet and vortexed immediately until the pellet was completely dissolved and the color was changed into olive green so that all the protein components were degraded. The mixture was then incubated in a water bath or heat block at 65℃ for 5 minutes. After incubation, 100 µl of isopropanol (100%) was added and mixed by inversion until DNA was precipitated as visible threads. The tube was centrifuged for 5 minutes at 10000×g. The supernatant was discarded and the pellet was dried by keeping the tube inverted state on a clean tissue paper for one minute. 100 µl of 70% ethanol was added and vortexed for 5 seconds. The tube was centrifuged for 5 minutes at 10000×g. The supernatant was carefully aspirated using a micropipette and keeping the micro-centrifuge tube in the inverted state on the tissue paper to allow the pellet to air dry for at least 5 minutes. Over-drying was avoided as the process can make it difficult to dissolve the DNA. Depending on the pellet size, 25-50 µl of nuclease free water was added and the tube was vortexed for 5 seconds and the mixture was incubated at 65°C for one hour in water bath for dissolving DNA or left overnight at room temperature. Nuclease free water was used instead of FG3 buffer (Elution buffer). Finally, the concentration and the purity of the DNA was measured using a Nano drop machine and adjusted the concentration for PCR.

#### 2.3.2. Polymerase Chain Reaction (PCR) Amplification of TSHR Gene

The isolated DNA was then amplified by PCR using TSHR gene-specific primers. At first, we performed PCR using primers set that could flank the sequence between Exon 1 to Exon 10, since global data showed that most of the common mutations in the TSHR gene of the patients with Congenital Hypothyroidism were confined in this region. Next, we conducted PCR for other regions of the TSHR gene. The primer sequences are listed in the **Table 1** as follows. To amplify the desired target sequence of TSHR gene, PCR amplification was conducted on a thermal cycler (Bio-Rad, USA). The final reaction volume was 10 µl for each of the reactions which contained 1 µL 10X PCR buffer, 0.3 µL 25mM MgCl2, 2 µL 5X Q-solution, 1.6 µL 2.5 mM dNTPs mixture, 0.2 µL 10mM Forward primer and 0.2 µL Reverse primer, 0.05 µL Taq DNA Polymerase, 50 ng of genomic DNA and total reaction volume was made up to 10µL by addition of nuclease free water. The thermal cycling condition included (a) initial denaturation at 95°C for 5 minutes, (b) cyclic denaturation at 95°C for 40 seconds and annealing at 58°C for 35 seconds and extension at 72°C for 40 seconds; and (c) final extension at 72°C for 5 minutes for 35 cycles.

**Table 1.**
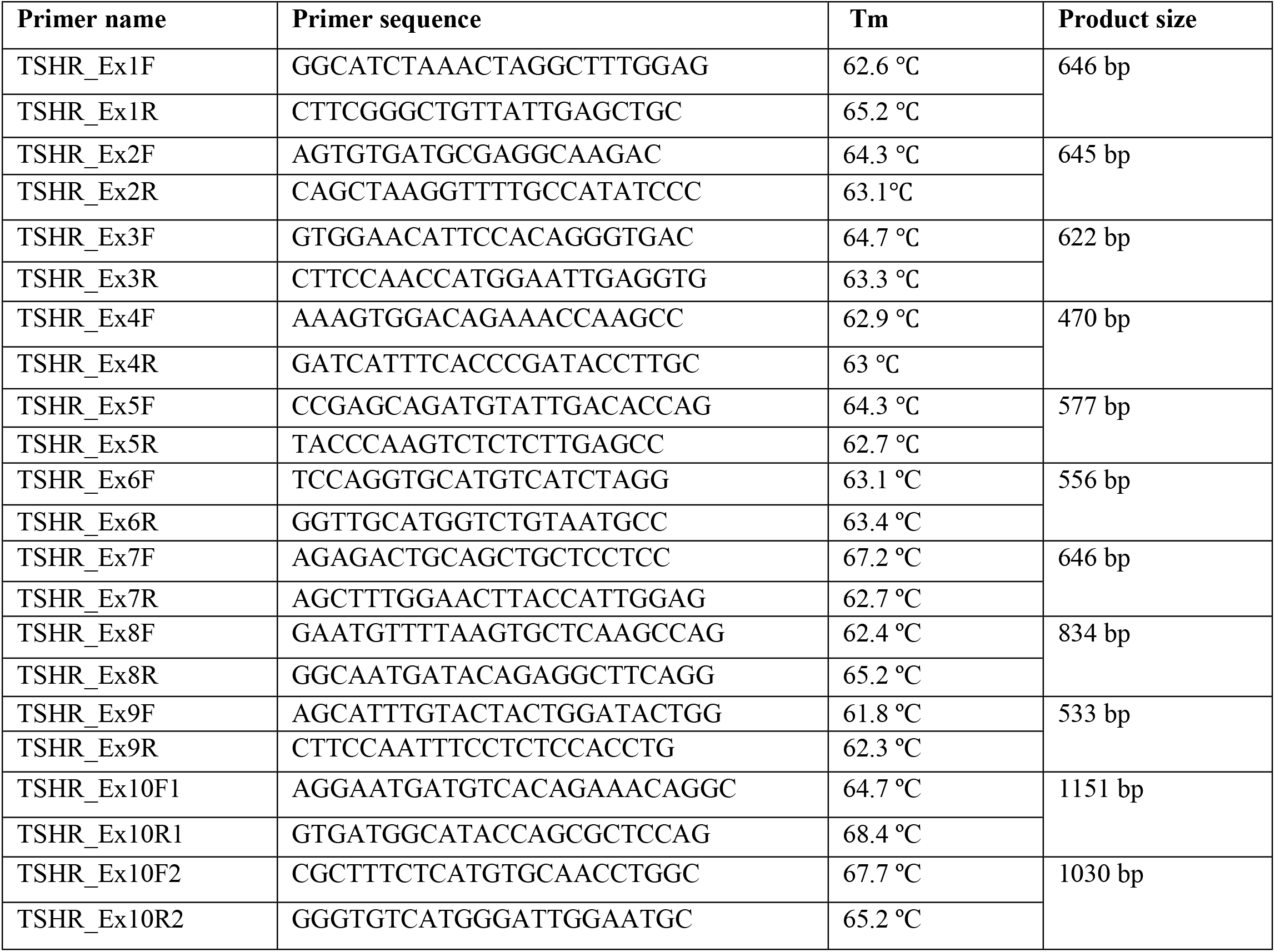
List of primers for PCR amplification and Sanger sequencing of TSHR gene.

### 2.4. Effect of Mutation on Predicted 3D Structure of Transmembrane Region of TSHR Protein

After performing the Sanger Sequencing technique, we detected 2 mutations in Transmembrane (TM)-region of TSHR protein. Since the small molecules drug binds to the TM region of TSHR protein, we targeted this region to see the effect of mutations in that particular site. TSHR protein is composed of a total of 764 amino acids and the TM region belongs to 368 to 764 amino acids of the full length TSHR protein. We used I-TASSER tool to predict the 3D structures of TM-region namely TSHR 368-764 by following the procedures as mentioned earlier for TPO protein. We selected two promising small molecules ligands (MS437 and MS 438) which act as agonist to TSHR protein. Finally, the molecular docking was performed for wild-type and mutant proteins using Autodock Vina protocol. Grid box center was x = 72.5922; y = 72.4245; z = 72.6927 and Grid box size was 25 in every axis during docking. Non bond interactions were also observed both for of MS437 and MS438 molecules.

### 2.5. Molecular Dynamics (MD) Simulation

The simulation was implemented through YASARA suits (9) employing AMBER14 force field (10) for calculations. The membrane was built during simulation. YASARA scanned the plausible transmembrane region comprising hydrophobic residues among the secondary structure elements of proteins (11). The protein was projected to the membrane, YASARA presented the recommended membrane insertion with required size (69.2Å∗ 7.3 Å) containing phosphatidyl-ethanolamine, -choline, and -serine lipid constituents. The whole simulation system was equilibrated for 250 ps. During the equilibration phase, membrane was artificially stabilized. The entire environment was equilibrated at 310K temperature with 0.9% NaCl and water solvent. The temperature was controlled by Berendsen thermostat process during simulation. The particle Mesh Ewald algorithm maintained long-range electrostatic interactions. The periodic boundary condition was applied for the whole simulation. The time step was 1.25 fs during 50 ns MD simulation. The snapshots were collected at every 100 ps. Diverse data containing root mean square deviation (RMSD), root mean square fluctuation (RMSF), total number of hydrogen bonds, radius of gyration, solvent-accessible surface area (SASA), and molecular surface area (MolSA) were retrieved from MD simulations, following our earlier MD data analysis (12).

### 2.6. Principal Component Analysis (PCA)

MD simulation data were applied to explore the structural and energy variabilities via principal component analysis (PCA) among TSHR-small molecule ligand (MS437 and MS438) complexes. The different multivariate energy factors were employed to explore the existing variability in the MD trajectory applying the low-dimensional space (13). The variables from MD data were bond distances, bond angles, dihedral angles, planarity, van der Waals energies, and electrostatic energies considered for the structural and energy factors (14). The data pre-processing were implemented by centering and scaling. In the analysis, final 45 ns MD trajectories were applied for exploration of the variations. The PCA model is reproduced by the following equation:

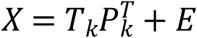

Where, multivariate factors are presented into the resultant of two new matrices by *X* matrix i.e. *T*_*k*_ and *P*_*k*_; *T*_*k*_ matrix of scores correlates the samples; *P*_*k*_ matrix of loadings associates the variables, *k* is the number of factors available in the model and *E* demonstrates the matrix of residuals. The trajectories were analysed through R, RStudio and essential codes. The R package ggplot2 was utilized for PCA plots generation.

## 3. Result

### 3.1. Investigation of Mutation in TSHR Gene

All the 21 patients with Dysgenesis had mutation in Exon 10 among a total of 11 Exons in TSHR gene. The mutations we found namely, c.1523C>T (p.Ser508Leu) and c.2181G>C (p.Asp727Glu). Among the 21 patients, only one patient had mutation c.1523C>T (p.Ser508Leu) and 20 patients had the other variant c.2181G>C (p.Asp727Glu). The variants were then analyzed by bioinformatics tools to explore the pathogenic effect. Firstly, the mutations were tested by Polyphen 2, Mutation taster and PROVEAN bioinformatics tools to see whether they possessed damaging effect or not. We found that the mutation c.1523C>T probably had damaging effect and c.2181G>C variant showed benign effect. **Fig. 1** represents a chromatogram showing the mutation (c.1523C>T) for the specific participant and **Table 2** shows the mutation found in TSHR gene.

**Table 2.** Mutation detection in the TSHR gene of hypothyroid patients and analysis of the effect of mutation using different bioinformatics tools.

| Nucleotide position | Amino acid position | Polyphen 2 | Mutation Taster | PROVEAN |
| --- | --- | --- | --- | --- |
| c.1523C>T | p.Ser508Leu | Probably Damaging | Disease causing | Deleterious |
| c.2181G>C | p.Asp727Glu | Benign | Benign | Neutral |

**Fig. 1.**
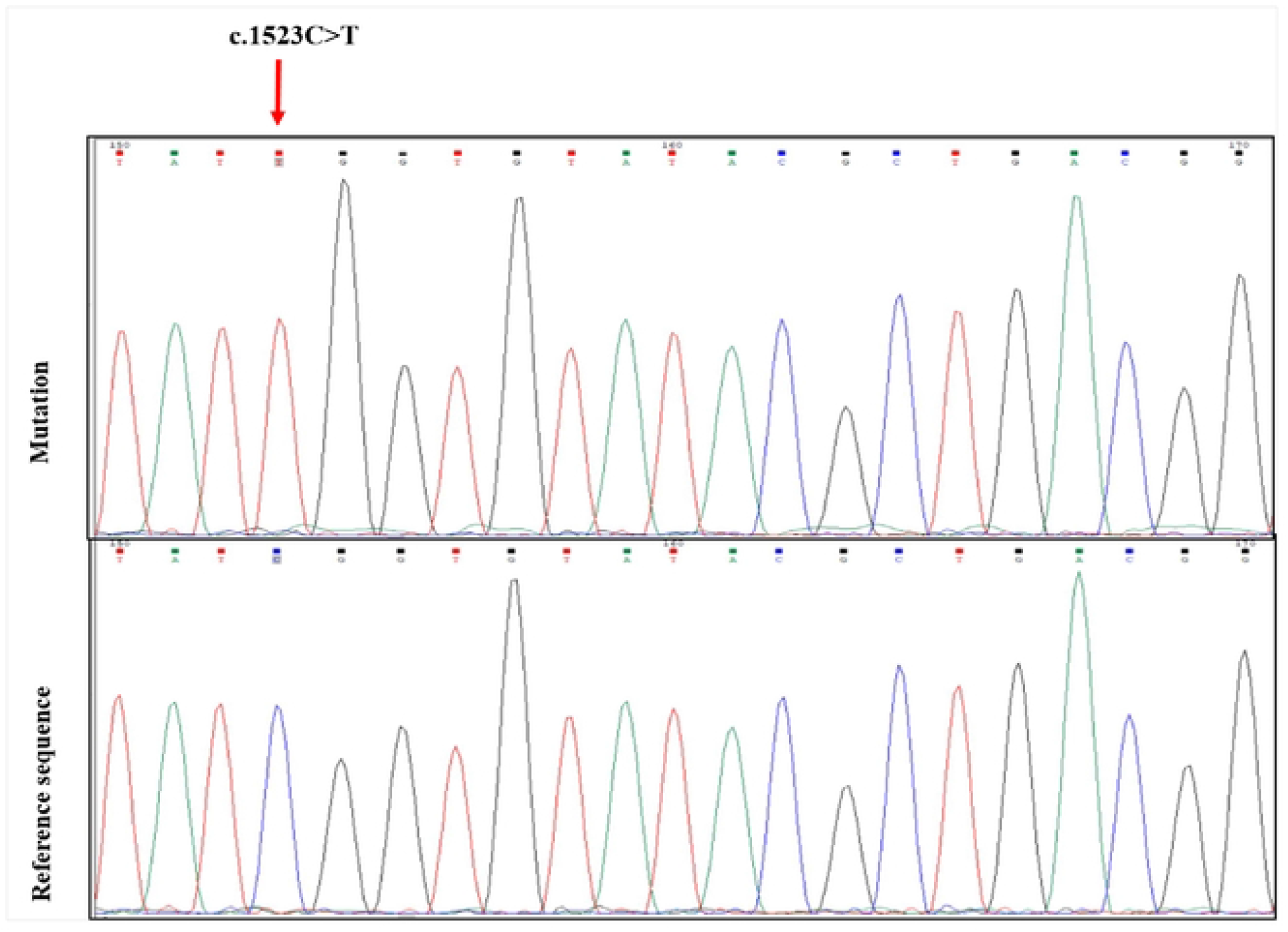
A representative chromatogram showing a mutation in Exon 10 of TSHR gene.

### 3.2. Modelling of the Transmembrane Region of TSHR Protein

Since the small molecules drug bind to the transmembrane region of the TSHR protein, we predicted the structure by using I-TASSER tool according to previously described TPO protein. The predicted structures were designated as wild type TSHR _**368-764**_ WT, p.Ser508Leu variant as TSHR _**368-764**_ MT1 and p.Asp727Glu variant as TSHR _**368-764**_ MT2. **Table 3** shows the C-score for wild type, MT1 and MT2 was −0.71, −0.61 and −0.63, respectively. The TM score and RMSD were 0.62±0.14, 8.4±4.5Å for TSHR _**368-764**_ WT; 0.64±0.13 and 8.2±4.5Å TSHR _**368-764**_ MT1; 0.63±0.13 and 8.2±4.5Å for TSHR _**368-764**_ MT2. **Fig. 2** depicts the 3D structures of the TM-region of TSHR protein and the small molecules MS437 and MS438.

**Table 3.** Summary of the corresponding model numbers, C–score, TM–score and the RMSD–score of the predicted 3D structures of TSHR 368-764 WT, TSHR 368-764 MT1 and TSHR 368-764 MT2.

| Features | TSHR 368-764 WT | TSHR 368-764 MT1 | TSHR 368-764 MT2 |
| --- | --- | --- | --- |
| Model no. | 01 | 01 | 01 |
| C-score | -0.71 | -0.61 | -0.63 |
| TM-score | 0.62±0.14 | 0.64±0.13 | 0.63±0.13 |
| RMSD (Å) | 8.4±4.5Å | 8.2±4.5Å | 8.2±4.5Å |

C-score=Confidence score range: [-5, 2]; TM-score=Template Modelling score, TM-score < 0.17 indicates random similarity and TM-score > 0.5 indicates correct similarity; RMSD=Root Mean Square Deviation. WT = Wild type, MT1 = Mutant 1 (p.Ser508Leu) and MT2 = Mutant 2 (p.Asp727Glu).

**Fig. 2.**
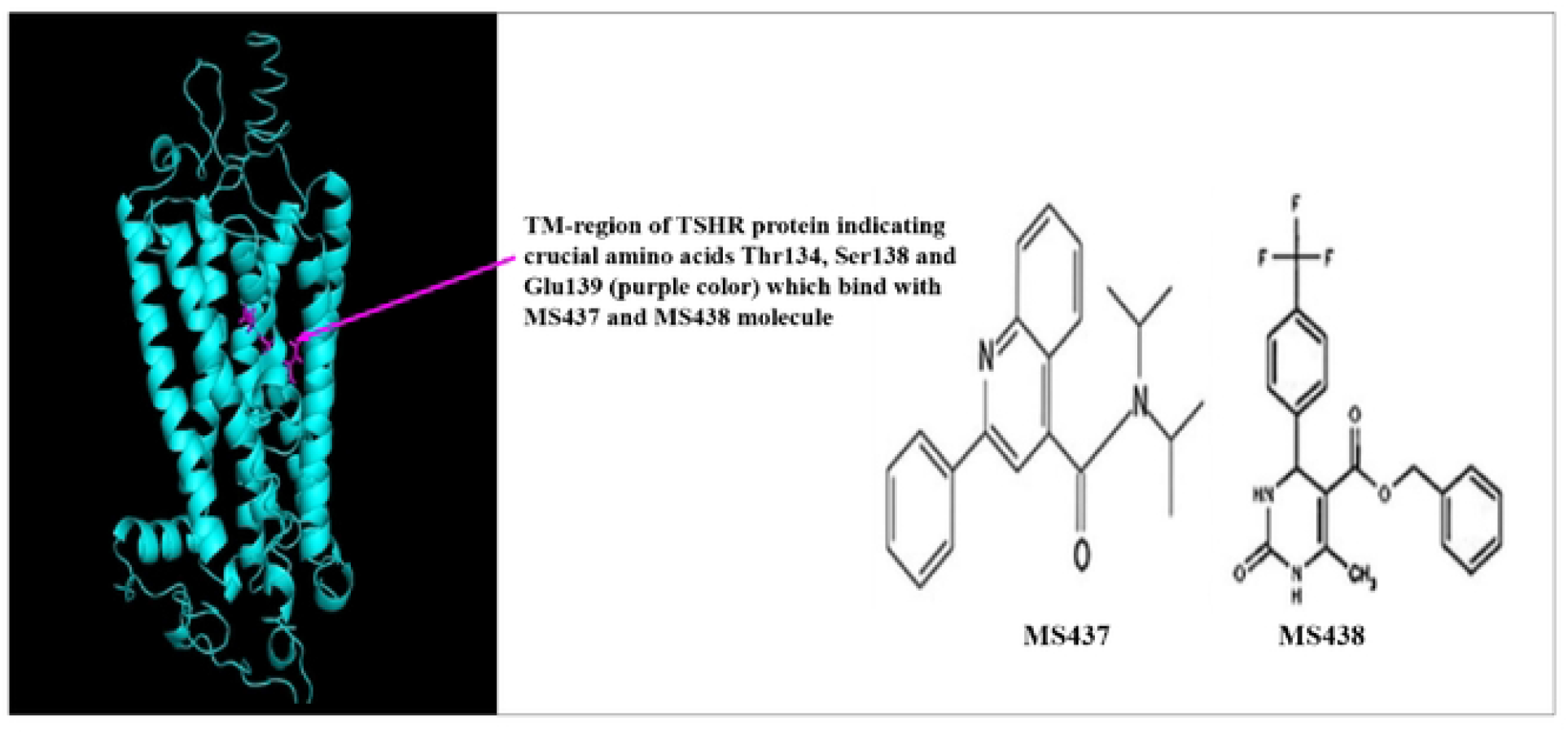
The predicted 3D structures of transmembrane region of TSHR proteins (368-764) showing crucial amino acids responsible for binding with MS437 and MS438 molecules.

### 3.3. Molecular Docking of MS437 and MS438 with TM Region of TSHR Proteins (Wild Type and Mutant)

The structures of the small molecules MS437 and MS438 were optimized and molecular docking was performed using PyRx software. The **Table 4** showed that the binding affinity of the wild type TSHR protein (−6, kcal/mol, TSHR_368-764_WT) was higher compared to the mutant cases (−4.8 kcal/mol for TSHR_368-764_ MT1 and −5.7 kcal/mol for TSHR_368-764_ MT2). Total non-bond interactions were 11, 19 and 12 for wild type, MT1 and MT2, respectively. MS437 binds to 501 Threonine and MS438 binds to Serine 505 and Glutamic acid 506 of transmembrane helix3 (TMH3) in full length TSHR protein with corresponding amino acid position Thr134, Ser138 and Glu139, respectively in TM-region (15). We tried to investigate whether these crucial amino acids could interact with small molecules thyrogenic drug. We found that in case of MS437 none of these three amino acids could interact with both wild type and mutant cases. On the other hand, MS438 interacted with all the crucial amino acids including Thr134, Ser138 and Glu139 for wild type case and for the mutant cases (TSHR_368-764_ MT1 and TSHR_368-764_ MT2), it could interact only with Ser138. The binding affinities were −7.1 kcal/mol, −5.4 kcal/mol and −2.6 kcal/mol for TSHR_368-764_ WT, TSHR_368-764_ MT1 and TSHR MT2, respectively. In **Table 5** the binding affinity of wild type protein was much higher for both of the ligands (MS437 and MS438). All the non-bond interactions were depicted in **Fig. 3** and **Fig. 4** for MS437 and MS438 respectively.

**Table 4.**
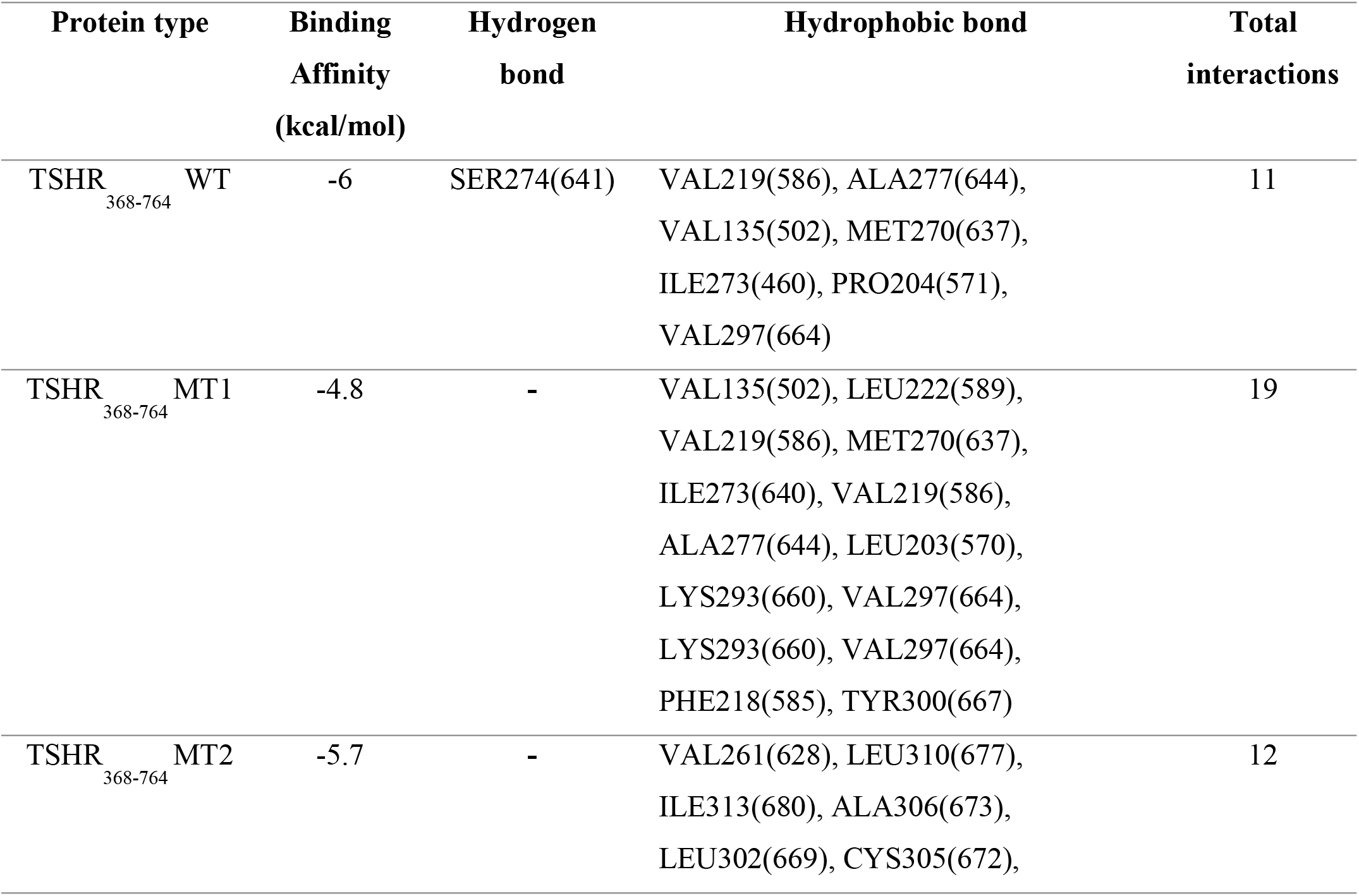
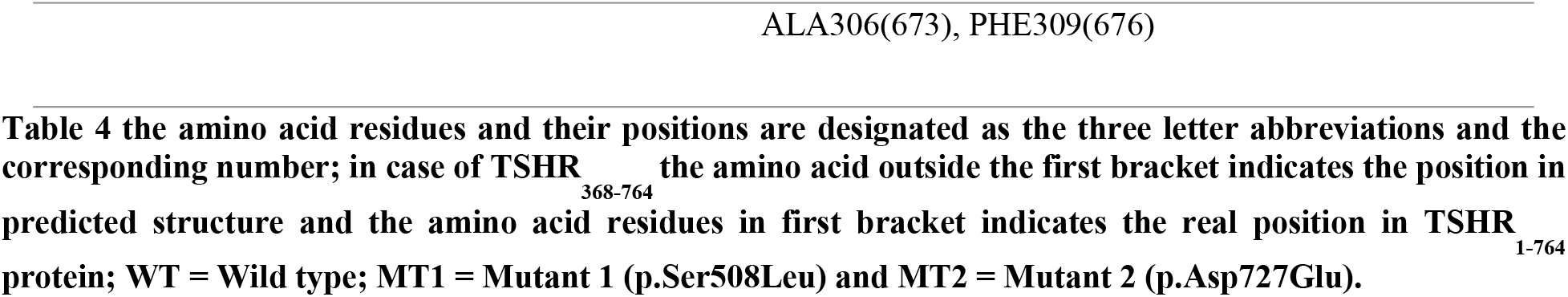
Binding Affinity and non-bond interactions of MS437 with TSHR368-764WT, TSHR368-764 MT1 and TSHR368-764 MT2 proteins after flexible docking.

**Table 5.** Binding Affinity and non-bond interactions of MS438 with TSHR368-764WT, TSHR368-764 MT1 and TSHR368-764 MT2 proteins after flexible docking.

| Protein type | Binding Affinity (kcal/mol) | Hydrogen bond | Electrostatic bond | Hydrophobic bond | Total interactions |
| --- | --- | --- | --- | --- | --- |
| TSHR<sub>368-764</sub> WT | -7.1 | ASN223(590),<br>VAL135(502),<br>SER138(505),<br>GLU139(506),<br>ASN223(590), | - | THR134(501),<br>VAL142(509),<br>MET270(637),<br>VAL297(664) | 13 |
| TSHR<sub>368-764</sub> MT1 | -5.4 | SER138(505),<br>ILE273(640),<br>VAL135(502),<br>SER274(641),<br>CYS202(569), | LYS293(660) | LEU100(467),<br>LEU203(570),<br>ALA277(640),<br>LYS293(660),<br>VAL297(664),<br>ILE273(640) | 14 |
| TSHR<sub>368-764</sub> MT2 | -2.6 | SER138(505),<br>LEU278(645),<br>SER274(641),<br>ALA277(644), | - | VAL219(586),<br>ALA277(644),<br>LEU278(645),<br>LEU100(467),<br>VAL297(664),<br>PRO301(668),<br>VAL219(586) | 14 |

**Fig. 3.**
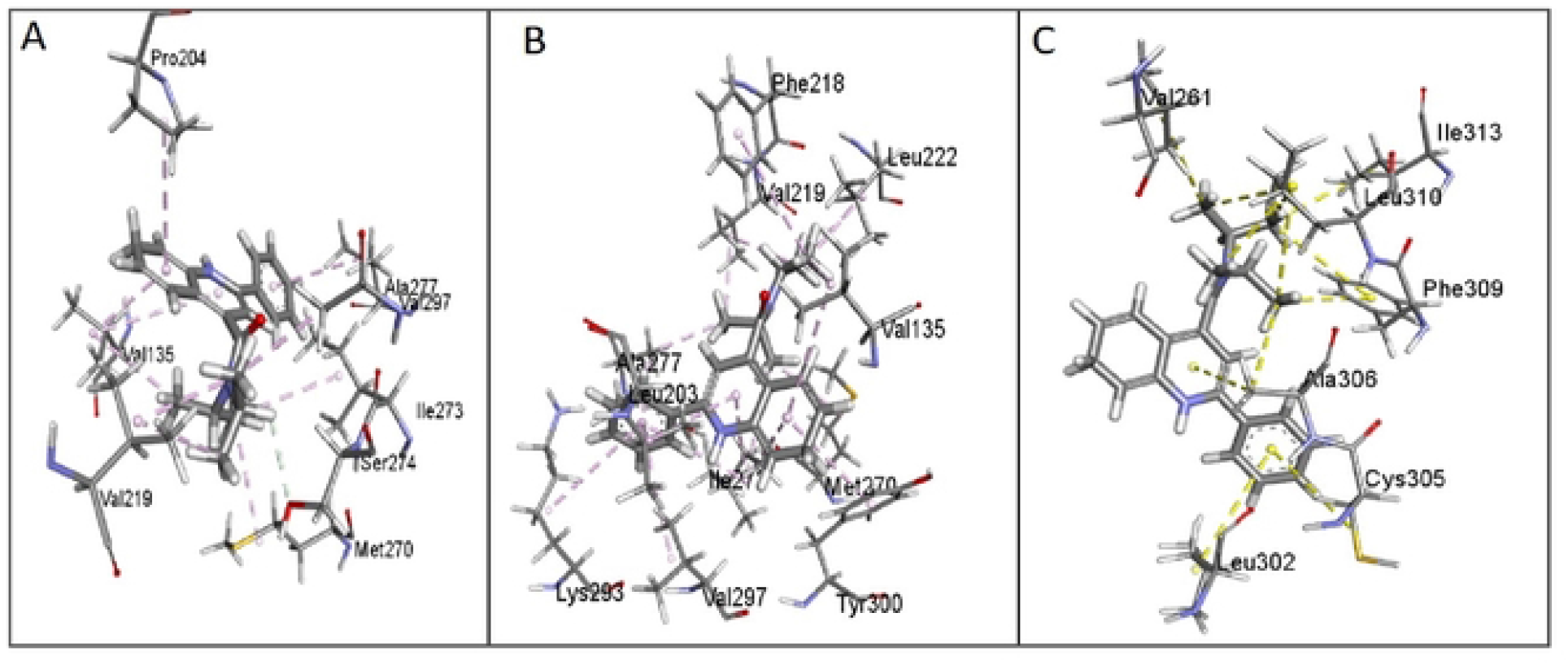
Non-bond interactions of MS437 with corresponding predicted structures as obtained using a BIOVIA Discovery Studio 2017. (A) TSHR368-764WT and MS437, (B) TSHR368-764MT1 and MS437 and (C) TSHR368-764MT2 and MS437.

**Fig. 4.**
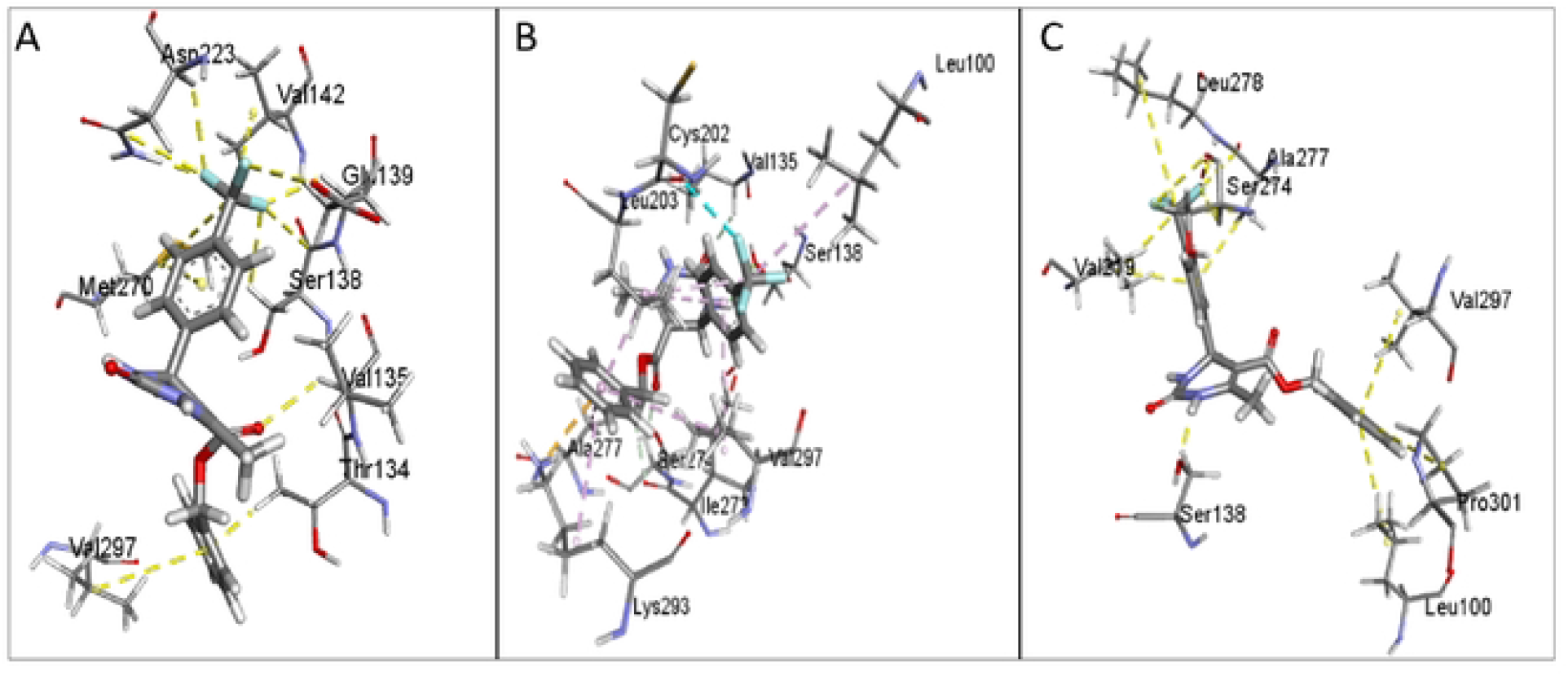
Non-bond interactions of MS438 with corresponding predicted structures as obtained using a BIOVIA Discovery Studio 2017. A. TSHR368-764WT and MS438 B. TSHR368-764MT1 and MS438 and C. TSHR368-764MT2 and MS438.

### 3.4. Molecular Dynamics Simulation

MD simulation has been implemented for each complex of TSHR_368-764_WT, TSHR_368-764_MT1, and TSHR_368-764_MT2 with two designated drugs (MS437 and MS438) for 50 ns time range.

In case of MS437 **(Fig. 5)**, the RMSDs for TSHR_368-764_MT2 (0.973-4.965 Å) displays less fluctuations for α-carbon atoms than TSHR_368-764_WT (0.906-5.91 Å), and TSHR_368-764_MT1 (0.939-5.504 Å) in **Fig. 5A**. Thus, suggesting that, comparatively MS437-TSHR_368-764_MT2 is stable in physiological conditions, while more fluctuation is visible in TSHR_368-764_WT at 43.8 ns (RMSD 5.91 Å) and TSHR_368-764_MT1 till 26.5 ns (RMSD ⁓5.2 Å).

**Fig. 5.**
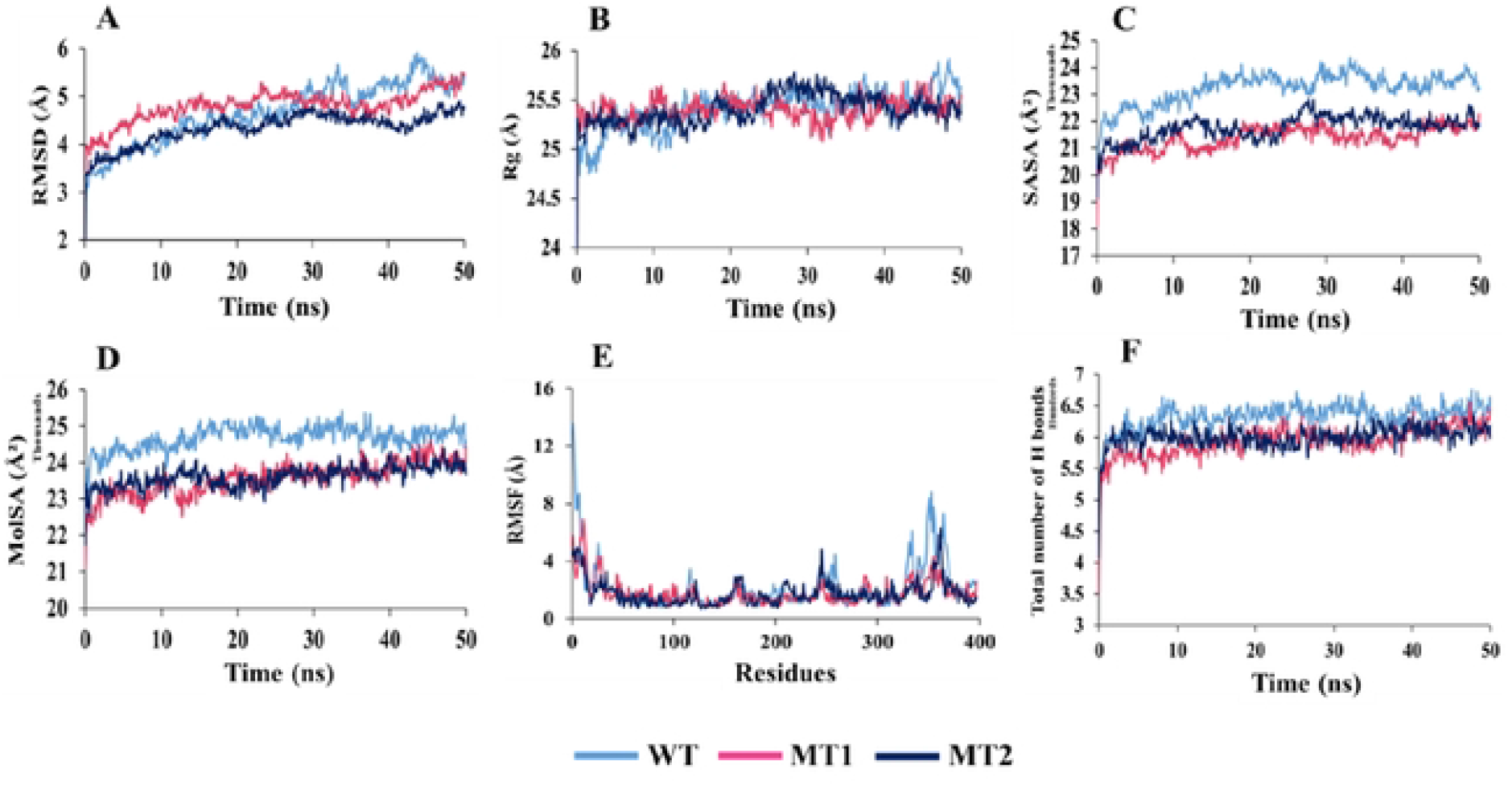
Analysis of 50 ns MD simulation of TSHR_368-764_ in complex with MS437 ligand. (A) Root mean square deviation values of C-α atom. The structural changes of TSHR_368-764_ proteins by means of (B) radius of gy ration. (C) solvent accessible surface area. (D) molecular surface area. (E) root means square fluctuations, and (F) total number of hydrogen bonds formed during the simulation.

The Rg manifests quite similar pattern in TSHR_368-764_MT1 and TSHR_368-764_MT2. However, TSHR_368-764_WT exhibits more fluctuations initially up to 5.5 ns and later from 46 ns. The average value remains ⁓25.40 Å for the three protein complexes. However, low compactness in ligand-mutant complexes is observed during simulation **(Fig. 5B)**.

In case of SASA, more deviations are found in TSHR_368-764_WT (18587.716-24360.945 Å^2^) compared to TSHR_368-764_MT1 (18106.599-22314.102 Å^2^) and TSHR_368-764_MT2 (19148.648-22851.2 Å^2^) complexes. However, TSHR_368-764_MT1 and TSHR_368-764_MT2 manifest some deviations at 7-19 ns, and 24-43 ns during simulation. Overall, mutant structures TSHR_368-764_MT1 and TSHR_368-764_MT2 are more stable as MS437 bound complexes than TSHR_368-764_WT **(Fig. 5C)**.

For MolSA in **Fig. 5D**, TSHR_368-764_WT shows (21208.043-25437.671 Å^2^) much fluctuations in the whole run than TSHR_368-764_MT1 (21135.321-24580.781 Å^2^) and TSHR_368-764_MT2 (21739.906-24397.155Å^2^). That means, TSHR_368-764_WT is unstable, but TSHR_368-764_MT1 and TSHR_368-764_MT2 are more stable as the ligand bound complexes in physiological condition.

The RMSF value deviates most for TSHR_368-764_WT in between 1(368)-30(397) and 326(693)-397(764) residues, where, TSHR368-764MT1 exhibits more fluctuations in 1(368)-47(415) residues than TSHR_368-764_MT2. Overall, TSHR_368-764_MT2 is more stable as a complex than others due to least deviation in the whole run **(Fig. 5E)**.

The total number of hydrogen bonds indicates structural rigidity of protein. In case of TSHR_368-764_WT high frequency of hydrogen bonds (average ⁓633) is observed during interaction while TSHR_368-764_-MT1 exhibits average 593, and TSHR_368-764_-MT2 manifests average 600 hydrogen bonds. Among the mutant structures, TSHR_368-764_-MT2 displays more structural stability in the simulation **(Fig. 5F)**.

During simulation, for MS438 **(Fig. 6)**, the RMSD values of α-carbon atoms remain ⁓5.251 Å in TSHR_368-764_-WT, ⁓5.53 Å in TSHR_368-764_MT1, and ⁓5.39 Å in TSHR_368-764_MT2. The fluctuations have been observed in TSHR_368-764_MT1 and TSHR_368-764_MT2 during 30-50 ns while least is found in TSHR_368-764_-WT. However, the average RMSD values of all the complexes are almost close **(Fig. 6A)**.

**Fig. 6.**
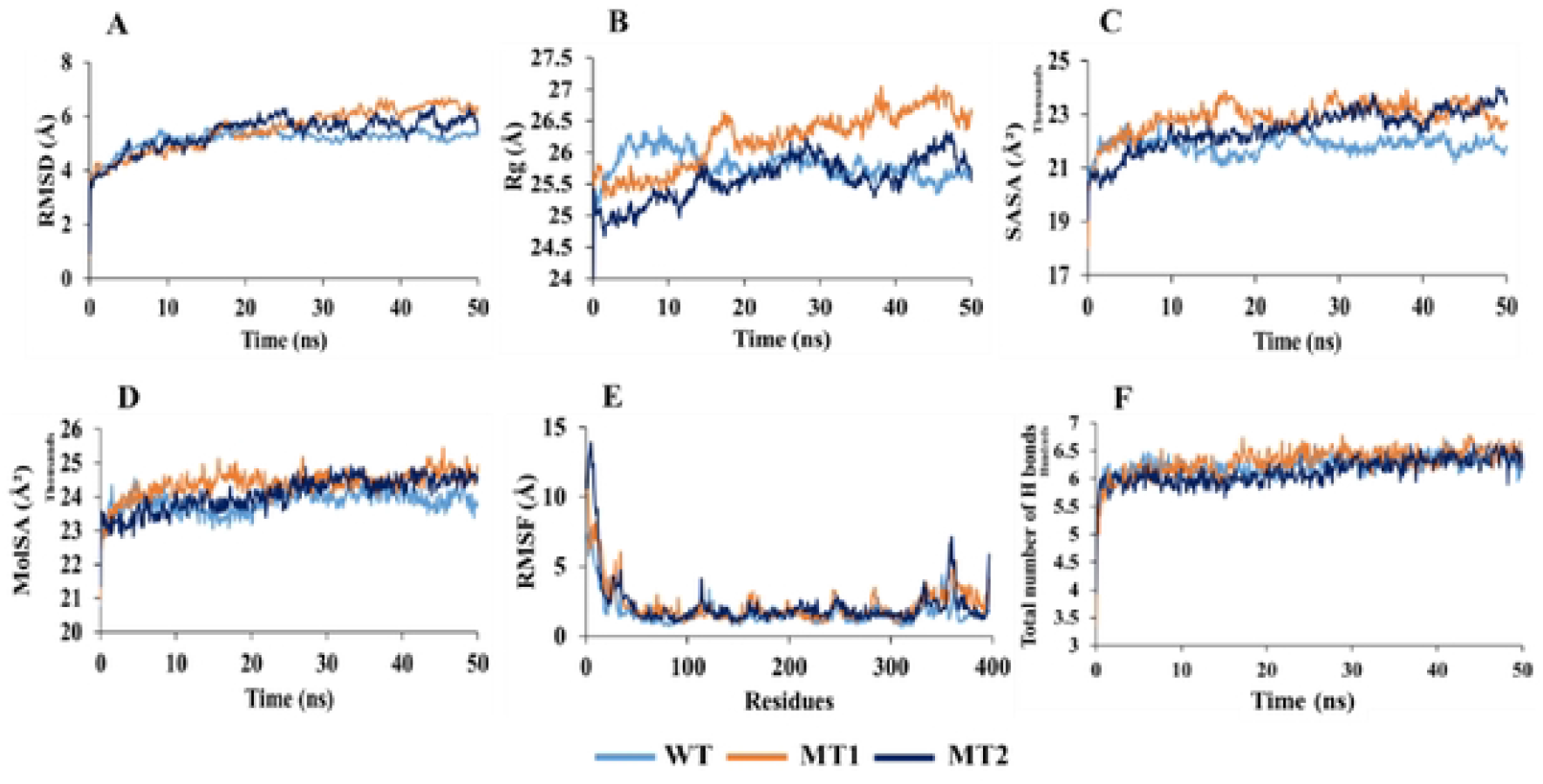
Analysis of 50 ns MD simulation of TSHR_368-764_ in complex with MS438 ligand. (A) Root mean square deviation values of C-α atom. The stnictural changes of TSHR_368-764_ proteins by means of (B) radius of gyration. (C) solvent accessible surface area. (D) molecular surface area. (E) root means square fluctuations, and (F) total number of hydrogen bonds formed during the simulation.

The Rg manifest quite high deviations among three complexes. However, TSHR_368-764_MT1 exhibits maximum 27.068 Å, which indicates higher stability than other complexes **(Fig. 6B)**. The SASA values remain close among three complexes. However, TSHR_368-764_-WT manifests least deviations during 10-20 ns and 30-50 ns **(Fig. 6C)**. In case of MolSA, the graphical pattern for three complexes are almost same through the whole MD run **(Fig. 6D)**.

The RMSF value more diverge in TSHR_368-764_MT2 till first 15 residues while least deviation is observed for TSHR_368-764_-WT through whole run. However, three complexes show quite similar pattern between 130(498)-240(608) residues **(Fig. 6E)**.

The highest number of hydrogen bonds (about 678) is observed for TSHR_368-764_MT1 while TSHR_368-764_-WT exhibit about 669 hydrogen bonds and TSHR_368-764_MT2 showed almost 665 to maintain stable conformation. Thus, TSHR_368-764_MT1 shows highest structural stability among the complexes **(Fig. 6F)**.

Moreover, we have visualized the binding pattern of MS437 and MS438 ligands with the wild type and mutants TSHR_368-764_ through the snapshots from MD simulation **(Fig. 7)**. In simulation, MS437 exhibits persistent interaction with the residues LEU302(669), ALA306(673), LEU310(677) of TSHR_368-764_MT2.

**Fig. 7.**
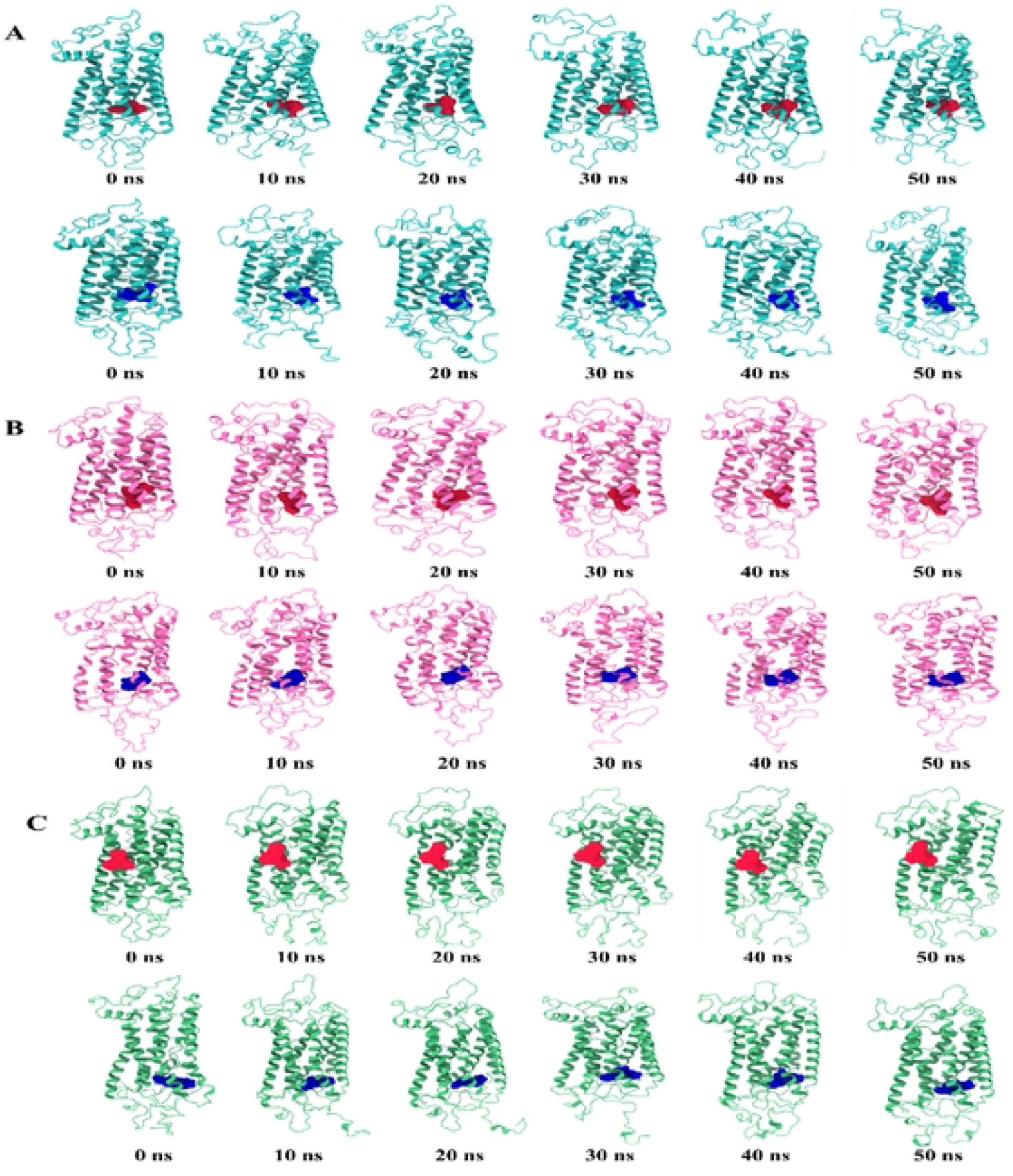
The snapshots of the generated conformers for TSHRα_368-764_ and ligands: MS437 (red). MS438 (blue) over the 50 ns MD simulation for (A) TSHR_368-764_-WT (cyan). (B) TSHR_368-764_-MTI (pink), and (C) TSHR_368-764_-MT2 (green) structures.

The MS438 ligand mostly shows stable interactions with the residues LEU100(467), VAL135(502), SER138(505), LEU203(570), PRO204(571), LYS293(660). Both ligands remain within the binding site in stable mutant proteins.

### 3.5. Principal Component Analysis (PCA)

Two PCA models are generated for structural and energy profiles of the protein-ligand complexes to assess and realize the dissimilarities among wild type and mutant proteins during MD simulation. The scores plot for MS437-protein **(Fig. 8A)** and MS438 protein **(Fig. 8C)** complexes have exhibited the different clusters for the wild type and mutants of TSHR_368-764_. It is observed in both protein-ligand complexes, TSHR_368-764_WT and TSHR_368-764_MT1 are remotely situated. Consequently, pathogenic TSHR_368-764_MT1 is liable for the differences. However, TSHR_368-764_WT and TSHR_368-764_MT2 are overlapped. The loading plots **(Fig. 8B and 8D)** demonstrate that bond, bond angles, van der Waals energies, and dihedral angles are closely distributed and displayed quite similar graphical patterns. This distribution mainly contributes for PC1 variance while coulomb energy difference contribute to PC2 variance. In MS437-protein complexes, the total 92.7% of the variance has been unveiled by PC1 and PC2, where PC1 expresses 76.1% and PC2 expresses 16.6% of the variance. Moreover, in MS438-protein complexes, the total 88.3% of the variance has been disclosed by PC1 and PC2, where PC1 expresses 72.1% and PC2 expresses 16.3% of the variance.

**Fig. 8.**
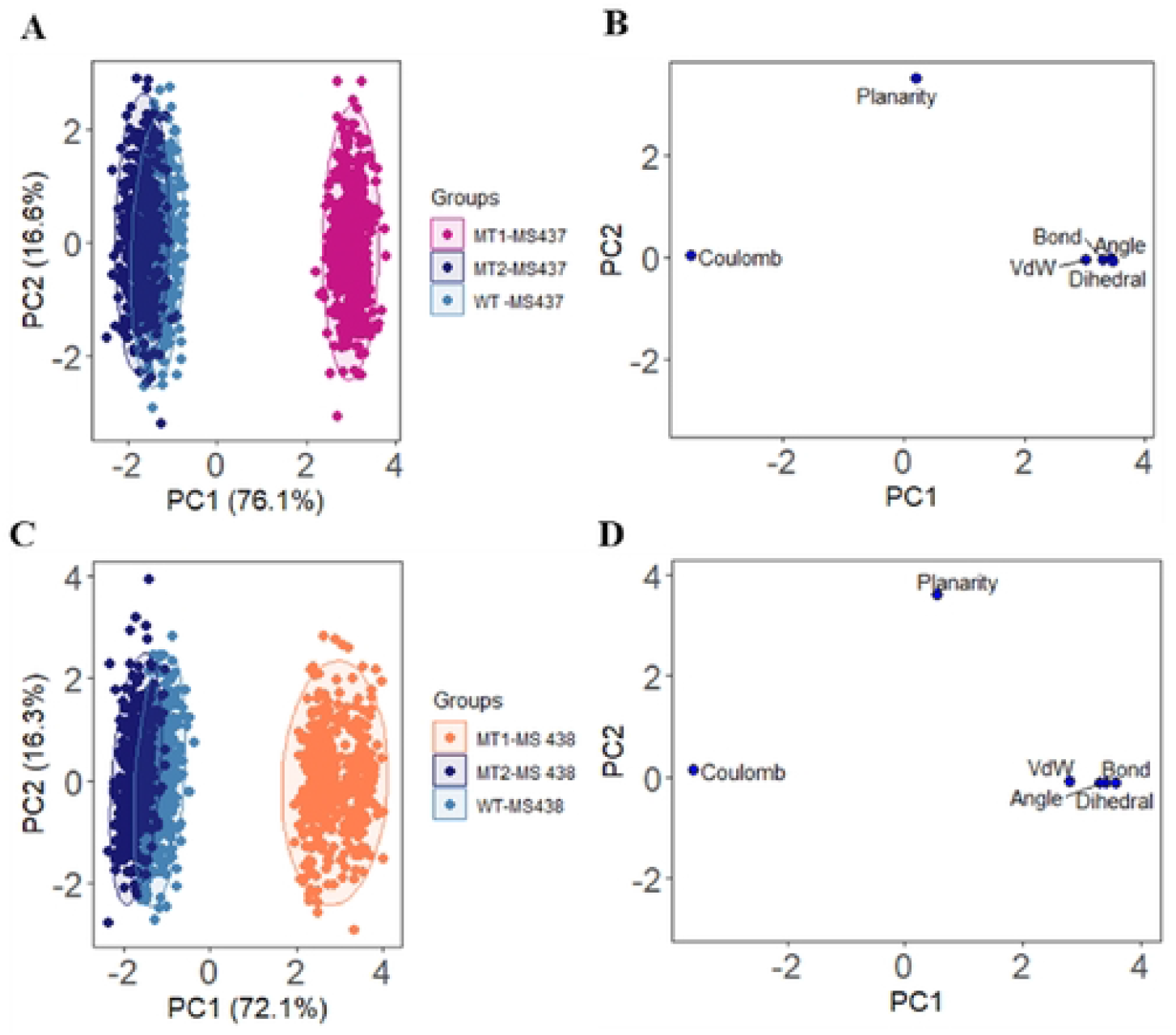
PCA analysis on 50 ns MD simulation. (A. C) The score plot represents three clusters for TSHR_368-764_ wild type and mutant protein structures, where each dot specifics one time point. The clustering is attributable as: WT-MS437 (sky blue). MT1-MS437 (pink). MT2-MS437 (navy blue), and WT-MS438 (sky blue). MT1-MS438 (orange). MT2-MS438 (navy blue). (B) Loading plot displays the energy and structural profile data from principal components analysis.

## 4. Discussion

The newborn screening for endocrine disorders is not frequently practiced in Bangladesh. In this study, we have focused on the etiology of dysgenesis types of CH patients having small glands or ectopic gland or agenesis (absent of thyroid gland). Different studies suggested that TSHR was the major gene responsible for growth and development of thyroid gland (2, 16). TSH binds to the receptor and creates the signaling pathway through G-protein coupled-receptor and Cyclic AMP-mediated adenylate cyclase. The full-length protein structure of TSHR is still under investigation through crystallography. Only a low small extracellular part of TSHR is described by crystallography. Mutations in the TSHR gene results from loss or gain of function of the protein that causes different phenotypic variations and lead to hyperthyrotopinemia to severe Congenital Hypothyroidism (17–19). Analysis of TSHR gene showed that two mutations were found, namely c.1523C>T and c.2181G>C in the patients and we analyzed the effect of mutations by using different bioinformatics tools. Almost all the tools were very much popular to analyze the mutational effect such as Polyphen 2, Mutation Taster and PROVEAN. The mutation c.1523C>T was found to be damaging, disease causing or deleterious and c.2181G>C was found to be benign or neutral. Since MS437 and MS438 are thyrogenic potent molecules, we selected these two molecules as ligands for molecular docking with the wild type and mutant structures of TSHR protein (15). The molecular docking analysis showed that the binding affinity for both of the ligands with mutant cases were decreased compared to the wild type TSHR protein. There is no published data based on c.1523C>T mutation but c.2181G>C was found as a normal variant. A study showed that although Asp727Glu was found as a normal variant it was also identified in follicular thyroid carcinomas (20). MD simulation indicates that, the RMSDs for MS437-TSHR_368-764_MT2 (average 4.37Å) shows less deviations for α-carbon atoms. Thus, proposing the complex as most stable in biological environments. However, MS438-protein complexes manifest quite close average RMSD values during simulation. The Rg of MS437-TSHR_368-764_WT exhibits more instabilities at start and end of the simulation. Conversely, TSHR_368-764_MT1 and TSHR_368-764_MT2 displays quite similar pattern of lesser compactness as well as more stability for interaction with MS437. In case of MS438, TSHR_368-764_MT1 exhibits highest Rg value, which identifies higher stability than other complexes. The analysis of SASA and MolSA values reveals that TSHR_368-764_MT1 and TSHR_368-764_MT2 mutant structures are more stable in their complex form with MS437 than TSHR_368-764_WT.

However, the SASA presents close pattern and MolSA exhibits almost same graphical pattern among the three complexes for the interaction with MS438.

In case of hydrogen bonds, MS437-TSHR_368-764_-MT2 manifests average 600 hydrogen bonds which is close to TSHR_368-764_WT (average ⁓633). The complex is more stable than the other mutant. On the other hand, MS438-TSHR_368-764_MT1 displays maximum structural stability compared to other complexes. Considering RMSF values, MS437 renders more stability to TSHR_368-764_MT2 than others. The RMSF value more diverges in TSHR_368-764_MT2 while minimum deviance is detected for TSHR_368-764_-WT and TSHR_368-764_MT1 mostly remains between both complexes. However, three complexes display almost similar stability between 130(498)-240(608) residues while interacting with MS438. Moreover, PCA analysis for MS437-protein and MS438-protein complexes have revealed the existing differences among structural and energy profiles of the structures. It is observable that TSHR_368-764_MT1 exhibits much variations than TSHR_368-764_WT and TSHR_368-764_MT2, emphasizing more damaging pattern in TSHR_368-764_MT1.

In this study, we have utilized allosteric ligands MS437 and MS438 as agonists against the identified mutants for TSHR_368-764_. These two ligands have ‘drug-likeness’ as well as previously confirmed their *in vivo* efficacy in animal studies (21). The agonists (MS437 and MS438) display different binding sites in the TMD of TSHR (15). After analysing all data, it can be proposed that low-affinity binding infers, a comparatively high concentration of the ligands can maximally occupy the binding sites to achieve maximum physiological response. Moreover, modifying chemical properties or ligands with novel scaffolds targeting signal-sensitive amino acids surrounding the allosteric binding sites might lead to design agonists with even higher efficiency to activate TSHR (22) (23).

## 5. Conclusion

The study investigated the molecular etiology of thyroid dysgenesis. Sequencing-based analysis detected two mutation (p.Ser508Leu, p.Asp727Glu) in transmembrane (TM)-region of TSHR gene. The effect of mutations on TM-region of TSHR protein was investigated targeting by small molecules drugs (MS437 and MS 438) via *in silico* approach using bioinformatics tools. The damaging effect in drug-protein complexes of mutants was revealed by molecular docking, non-covalent interaction, molecular dynamics simulation, and principle component analysis. The findings will be helpful to realize the molecular etiology of thyroid dysgenesis (TH) via exploring the mutational impact for TSHR protein and suggest more efficient treatment strategies.

## Acknowledgements

The authors are grateful to University Grants Commission for its generous support and also thankful to the institute for developing Science and Health initiatives (ideSHi), Bangladesh and Dr. Mohammad A. Halim, Division of Computer Aided Drug Design, The Red-Green Research Centre, Bangladesh. This study was completed under the support from the Higher Education Quality Enhancement Project of the University Grants Commission (UGC) of Bangladesh.

